# Quiescence Induction Triggers Acute Accumulation of DNA Double-Strand Breaks That Drive Quiescence Heterogeneity

**DOI:** 10.1101/2025.09.24.678365

**Authors:** Kotaro Fujimaki, Andrew Paek, Guang Yao

**Affiliations:** Department of Molecular and Cellular Biology, University of Arizona, Tucson, AZ, 85721, USA; Arizona Cancer Center, University of Arizona, Tucson, AZ 85719, USA; Department of Systems Biology, Harvard Medical School, Boston, MA, 02115, USA

## Abstract

DNA double-strand breaks (DSBs) represent one of the most serious threats to genome integrity. Endogenous DSBs chiefly arise from cell cycle activities such as DNA replication and division, while dormant, non-proliferative cells are generally considered protected from such damage. Here we report that induction of quiescence by growth restriction in human retinal pigment epithelial (RPE) cells unexpectedly leads to rapid accumulation of DSBs within hours, reaching levels that exceed those in continuously proliferating cells. These DSBs occur in a cell cycle stage-dependent manner, predominantly in cells past the restriction point when quiescence is induced. Mechanistically, this DSB accumulation results from continued cell cycle activity under growth restriction conditions, accompanied by downregulation of DSB repair genes, allowing these breaks to persist during quiescence. Cells that accumulate more DSBs during quiescence induction enter a deeper state of quiescence, requiring stronger growth stimulation for cell cycle re-entry. Notably, cell cycle re-entry depends on DSB repair mediated by DNA-PK, an intrinsically error-prone process. Our findings establish quiescence induction as a previously unrecognized source of latent genome instability, with implications for tissue maintenance and aging where transitions between quiescence and proliferation are critical.

## INTRODUCTION

DNA damage and the cell cycle are closely intertwined, with each affecting the other. Various cell cycle activities inherently generate endogenous DNA damage^1–3^. For instance, spontaneous replication fork collapse during DNA synthesis^1,4–6^, as well as increased metabolism that is associated with cell cycle progression^7^, can result in up to 50 DNA double-strand breaks (DSBs) per cell per day^8^. Conversely, DNA damage, including DSBs, delays or arrests cell cycle progression, depending on the severity of the damage^9,10^. In contrast to proliferating cells in the cell cycle, quiescent cells, due to the absence of replication and division, are considered protected from DSB formation^11,12^.

DNA damage response (DDR) in vertebrates is coordinated by the PI3K-like kinases, ATM, DNA-PK, and ATR^13^. In addressing DSBs, the most critical among various types of DNA lesions that affect genome stability, ATM recruits DDR factors via phosphorylating the histone H2AX, triggers DNA damage checkpoints, and promotes a low-error DSB repair pathway via homologous recombination (HR)^13,14^. Alternatively, DNA-PK promotes an error-prone DSB repair pathway, non-homologous end joining (NHEJ)^13^. ATR, though not activated by DSBs, prevents DSBs during S phase by limiting replication fork collapse^13^. Notably, the H2AX phosphorylation, which serves as a platform for DDR factor recruitment, relies on growth signals^15,16^, and both HR and NHEJ pathways—which are critical for DSB repair—are down-regulated under conditions that restrict cell growth^11,17–20^.

In growth restriction conditions—such as upon mitogen withdrawal or contact inhibition— E2F activators, transcription factors essential for DNA replication and cell cycle progression^21^, are inactivated, leading cells to enter quiescence^22^. If cells encounter growth restriction conditions before reaching the restriction point (RP), they withdraw from the cell cycle and enter quiescence without proceeding further; otherwise, cells complete the current cell cycle before entering quiescence^22^. In quiescence, cells should be protected from cell cycle-incurred DSB formation^11,12^. However, extended periods in quiescence—spanning years in some adult stem cells—can nevertheless lead to gradual DNA damage due to declined mitochondrial and autophagy functions and consequential accumulation of reactive oxygen species (ROS)^17,23–25^. Consistently, aged hematopoietic stem cells (HSCs) exhibit higher levels of DNA damage compared to younger HSCs^17,26,27^, and such accumulation of DNA damage is considered detrimental to cellular function^28^. However, the surprising finding that even young quiescent HSCs could display more DNA lesions than their proliferating counterparts^17^ suggests the existence of an age-independent, “fast” mechanism for DNA damage formation in quiescent cells, although the mechanistic nature remains elusive.

Here, we investigated the DSB formation during quiescence entry in human cells, which we hypothesized could underlie the "fast" mechanism accumulating DNA damage in quiescent cells described above. Using single cell imaging, we demonstrate that quiescence induction upon growth restriction conditions evoked rapid DSB accumulation. The degree of the evoked DSBs was associated with the prior cell cycle position before quiescence entry. Specifically, cells that had passed the RP resulted in a higher level of DSBs than cells before the RP due to continued cell cycle activities upon quiescence induction. Consequently, post-RP quiescent cells exhibited reduced potential for cell cycle reentry. Repair of DSBs was critical for quiescence exit, and this process was mediated by error-prone NHEJ factor DNA-PK. Our findings suggest that the period between quiescence induction and quiescence entry represents a critical window for acute DSB accumulation, leading to heterogeneity and mutagenic potential in quiescent cells.

## RESULTS

### Quiescence induction evokes DSBs

To determine whether quiescence induction triggers DSB formation, we subjected a retinal pigment epithelial (RPE) cell line—engineered with endogenously tagged PCNA-mRuby and p21-GFP fluorescent reporters^10^ —to growth deprivation (serum and L-Glutamine starvation; hereinafter starvation), and quantified DSBs using neutral comet assay^29^. In this system, quiescent (G0) cells are identified as PCNA_low_/p21_high_, while proliferating (G1-S-G2/M) cells are PCA_high_/p21_low_ (Fig. S1A). Starvation induced the vast majority of asynchronously growing RPE cells into quiescence within 2 days, as indicated by 95% PCNA_low_/p21_high_ cells (Fig. S1A, B). Remarkably, the DNA% in comet tails showed a significant increase within 3 hours of starvation (Fig. 1A). It continued to rise over time (Fig. 1A, B) but largely plateaued by the 2nd day (Fig. 1C), suggesting that DSB accumulates primarily during quiescence entry and slows down once cells enter quiescence. We also subjected RPE cells to continuous culture for 2 weeks to reach full contact inhibition. The induced quiescent cell population, which increased over time as contact inhibition became more stringent (Fig. S1C), exhibited correspondingly increased DNA% in comet tails (Fig. S1C, D). Together, these data establish that quiescence induction triggers DSB accumulation in RPE cells as they transition from proliferation to quiescence.

**Figure 1.**
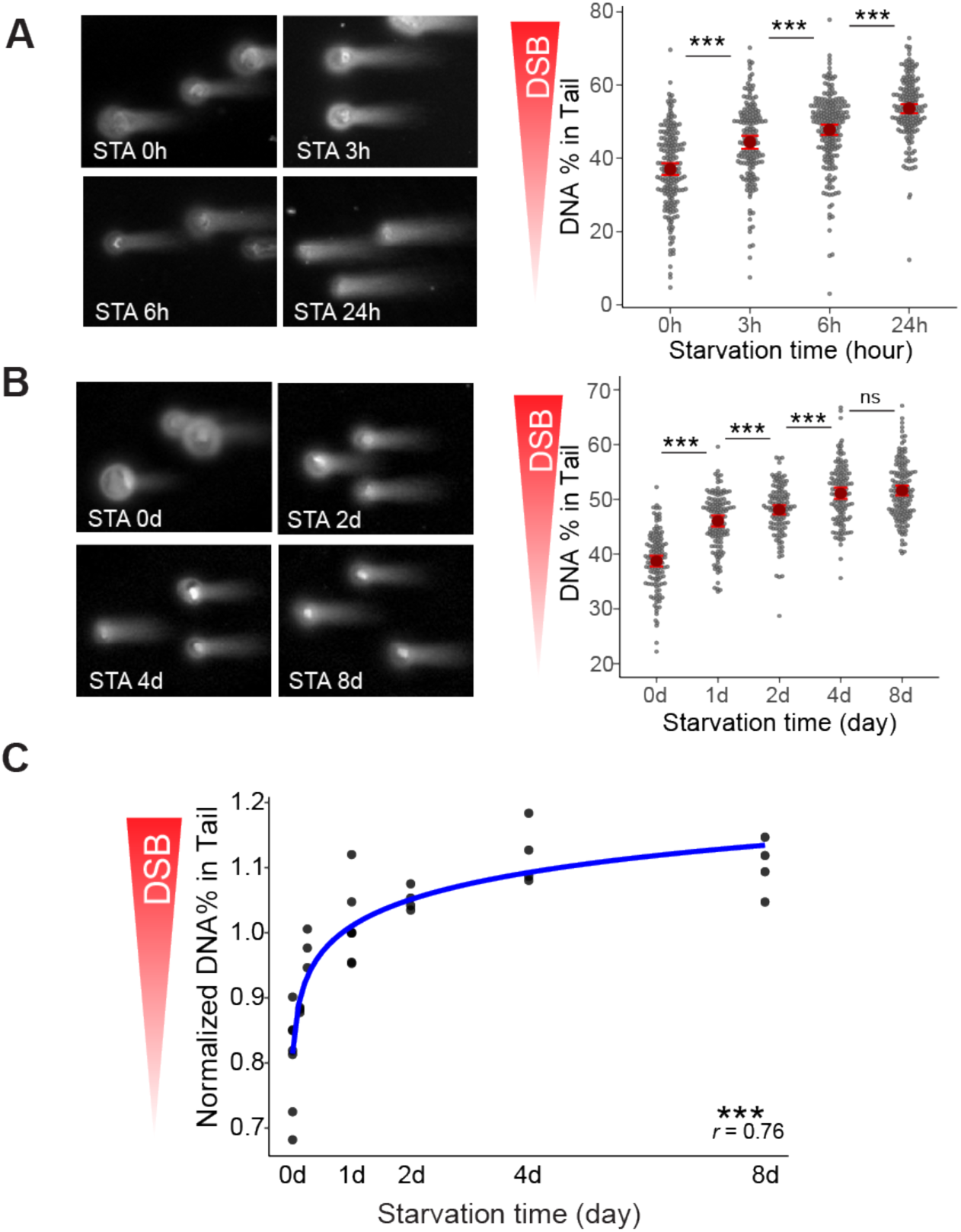
Quiescence induction evokes DSBs. (A-B) Cells, starved up to 24 hours (A) or 8 days (B), were analyzed by neutral comet assay. Left: representative neutral comets of cells upon starvation for the indicated time. Right: percent of DNA in the tail (n > 100 cells per condition). ns and *** indicate FDR-adjusted p > 0.05 and p < 0.001, respectively, in Mann Whitney U test. **(C)** Median percent of DNA in the tail (DSB index) of cells entering and maintaining quiescence from datasets A and B (n = 3 and 4 for A and B, respectively). Median DSB indices were normalized to 1d-starved cells. *** indicates the significant correlation (p < 0.001 in Spearman’s rank) between DSB indices and starvation time.

### Cell cycle activity under starvation leads to the accumulation of DSBs

Having established that quiescence induction triggers DSBs, we next sought to identify their sources. We focused on starvation, which induces quiescence more rapidly and consistently in the RPE cell population than contact inhibition. We observed that the expression of HR and NHEJ pathway genes decreased significantly following starvation (Fig. S2A), consistent with the expected downregulation of HR and NHEJ activities under growth restriction conditions^11,17–20^. We hypothesized that upon starvation, in cells past the RP, continued cell cycle activity—which is prone to causing damage—accompanied by down-regulated DSB repair leads to DSB accumulation (Fig. 2A).

**Figure 2:**
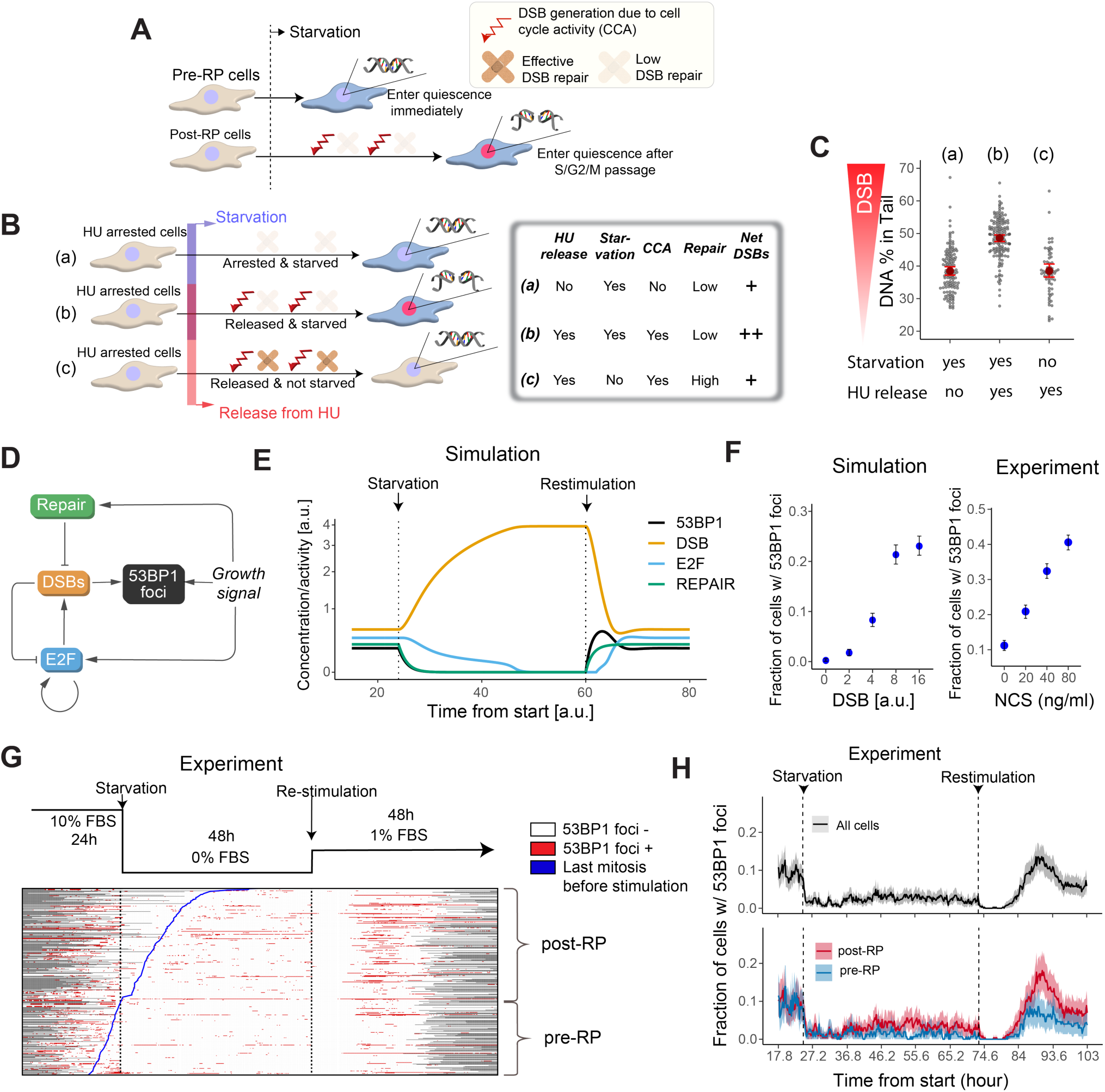
Cell cycle activity during starvation induces DSBs. **(A)** Proposed model for DSB generation. Cells induced into quiescence after passing the RP continue cell cycle activities under starvation, resulting in DSB accumulation due to damage-prone proliferation and reduced DNA repair. In contrast, cells induced into quiescence before the RP can enter quiescence with minimal cell cycle activity, avoiding DSBs. **(B)** Experimental scheme to test the model in (A). The same inset from (A) applies. The inset table summarizes experimental conditions and expected outcomes. **(C)** Percent of DNA in the comet tail of cells cultured under conditions (a), (b), and (c) from (B) (n > 60 cells per condition). **(D)** Circuit diagram of the components included in the mathematical model (see main text for details). **(E)** Simulated concentration/activity dynamics of components from (D) during quiescence entry and exit, shown as a result of deterministic simulation. **(F)** Left: The correlation between pre-stimulation DSB levels and the peak of the 53BP1 pulse in model simulation. The 53BP1 pulse peak reflects the maximum fraction of cells with 53BP1 foci post growth restimulation (n = 1700 simulated cells per condition; 53BP1 activity > 0.8 was considered foci positive). Right: Experimental equivalent (n > 1700 cells per condition), where RPE-53BP1 cells starved for 2 days were treated with indicated NCS concentrations for 1 hour and growth-stimulated for 18 hours before 53BP1 quantification. The 53BP1 pulse peak appears at ∼18 hours post-stimulation (as determined in G). Error bar, 95% confidence interval from bootstrapping. **(G)** 53BP1 foci dynamics during quiescence entry and exit (n > 380 cells). Experimental scheme shown above. In the heatmap, red and white indicate presence or absence of 53BP1 foci, respectively; gray, missing data. Cells are aligned by mitotic timing (blue line) before growth re-stimulation. Cells that did not divide during starvation period (i.e., no division between starvation and re-stimulation) were classified as pre-RP, while those that divided during starvation were classified as post-RP. (**H**) Fraction of cells with 53BP1 foci over time in pre- and post-RP groups – data from G (n > 150 and 230 cells for pre- and post-RP, respectively). Median and 95% confidence interval from bootstrapping are shown in thick lines and colored ribbons. Only time frames where more than 80% of all tracked cells have valid data are included.

To test whether the continued cell cycle activity under starvation induces DSBs, we first arrested cells at the G1/S boundary (beyond the RP) by hydroxyurea (HU, Fig. S2B) and then subjected them to three different conditions prior to DSB quantification using comet assay (Fig. 2B): (a) arrested & starved (i.e., cultured in starvation medium containing HU), (b) released & starved (in starvation medium without HU), or (c) released & not starved (in growing medium without HU). If cell cycle activity in starvation causes DSBs, the following is expected: in (a), arrested & starved, G1/S-arrested cells lack cell cycle activity and should not accumulate excess DSBs even under starvation; in (b), released & starved, cells resume cell cycle activity under starvation, which should result in excess DSBs; in (c), released & not starved, cells resume the cell cycle without starvation, thereby avoiding excess DSBs. Therefore, the expected DSB levels would be (a) < (b) and (b) > (c) (Fig. 2B). Indeed, this pattern was observed experimentally (Fig. 2C), suggesting that cell cycle activity under starvation induces DSBs.

### Quiescence induction leads to the accumulation of DSBs primarily in cells past the RP

To determine whether cell cycle position at the starvation onset affects subsequent DSB accumulation (Fig. 2A), we monitored the DSB reporter, 53BP1-Apple^30^ in starved cells. However, we initially observed that 53BP1-Apple underestimated DSB levels under starvation conditions— switching to starvation medium shortly before quantification decreased the fraction of cells positive for 53BP1 foci (Fig. S2D), despite the increase in DSBs following starvation (Fig. 1). This DSB underestimation by 53BP1-Apple was likely because growth restriction conditions attenuate phosphorylated H2AX (γH2AX)^15,16^, upon which 53BP1 recruitment depends.

To overcome this limitation and assess DSB levels in quiescent cells, we developed a coarse-grained mathematical model to identify an alternative index of DSB detection (Fig. 2D, Table S1, see *Methods* for detail). In this model, E2F transcriptional activators are stimulated by growth signals and promote cell cycle activities^21^, which subsequently generate DSBs. These DSBs promote 53BP1 recruitment and suppress cell cycle activities by inhibiting E2F^31^ until they are repaired by DNA repair machinery. DNA repair, 53BP1 recruitment, and E2F activity were modeled as dependent on growth signaling.

Model simulations indicated that although 53BP1 signal decreases during starvation even as DSBs accumulate, the true extent of DSBs in quiescent cells could be transiently revealed following growth restimulation. Specifically, growth restimulation permits 53BP1 recruitment to DSB sites. This effect is transient, as restimulation also activates DNA repair, which subsequently resolves DSBs, resulting in a characteristic pulse of 53BP1 signal (Fig. 2E) across a broad range of parameter combinations (Fig. S2E). The peak of the 53BP1 pulse, reflecting the fraction of cells exhibiting 53BP1 foci in model simulation, is positively correlated with the DSB levels before restimulation (Fig. 2F, left). These results suggest that growth restimulation provides a window during which pre-existing DSB levels can be revealed by the 53BP1 pulse following restimulation. To confirm that this 53BP1-pulse peak can be used to quantify pre-existing DSBs in quiescent cells, we treated quiescent cells with increasing doses of neocarzinostatin^32^ (NCS) to induce DSBs prior to growth restimulation—indeed, we observed a dose-dependent increase in the fraction of cells positive for 53BP1 foci (Fig. 2F, right).

Next, we experimentally monitored the 53BP1-Apple foci pulse in cells upon growth restimulation to determine the pre-existing DSB levels in these cells during starvation. The fraction of cells with 53BP1 foci dropped immediately after starvation and remained low for the 2-day starvation period (Fig. 2G, H, Fig. S2F, G), suggesting starvation-induced foci suppression despite ongoing DSB accumulation (Fig. 1C). Following growth restimulation, we observed a transient increase in 53BP1 foci, which peaked at around 18 hours post-stimulation and subsequently declined, showing a pulse-like pattern (Fig. 2H, Fig. S2G).

We classified RPE cells expressing 53BP1-Apple^30^ as either pre-RP (no division post-starvation) or post-RP (division post-starvation) at the initiation of starvation. Notably, the post-RP subset exhibited a higher fraction of cells positive for 53BP1 foci at 18-hour peak than the pre-RP subset (Fig. 2H, Fig. S2G). Together, our modeling and experimental results suggest that compared to cells prior to reaching the RP, cells past the RP upon starvation harbor more DSBs after entering quiescence (Fig. 2A).

### DSBs drive deep quiescence

Having observed that post-RP cells accumulate more DSBs during quiescence entry, we next asked whether this differential DSB accumulation has functional consequences. Cellular quiescence exists in a spectrum of depths: deeper quiescent cells require stronger growth stimulation to reenter the cell cycle than shallower counterparts^33–35^. Because stress and damage signals often increase quiescence depth^35–37^, we hypothesized that post-RP cells enter deeper quiescence than pre-RP cells due to excess DSBs accumulated. To test this hypothesis, we analyzed the proliferation tendency of quiescent RPE cells of pre- and post-RP origins. Upon growth restimulation, quiescent cells of the post-RP origin displayed a significantly delayed cell division compared to those of pre-RP origin (Fig. 3A and B), consistent with their deeper quiescent state.

**Figure 3:**
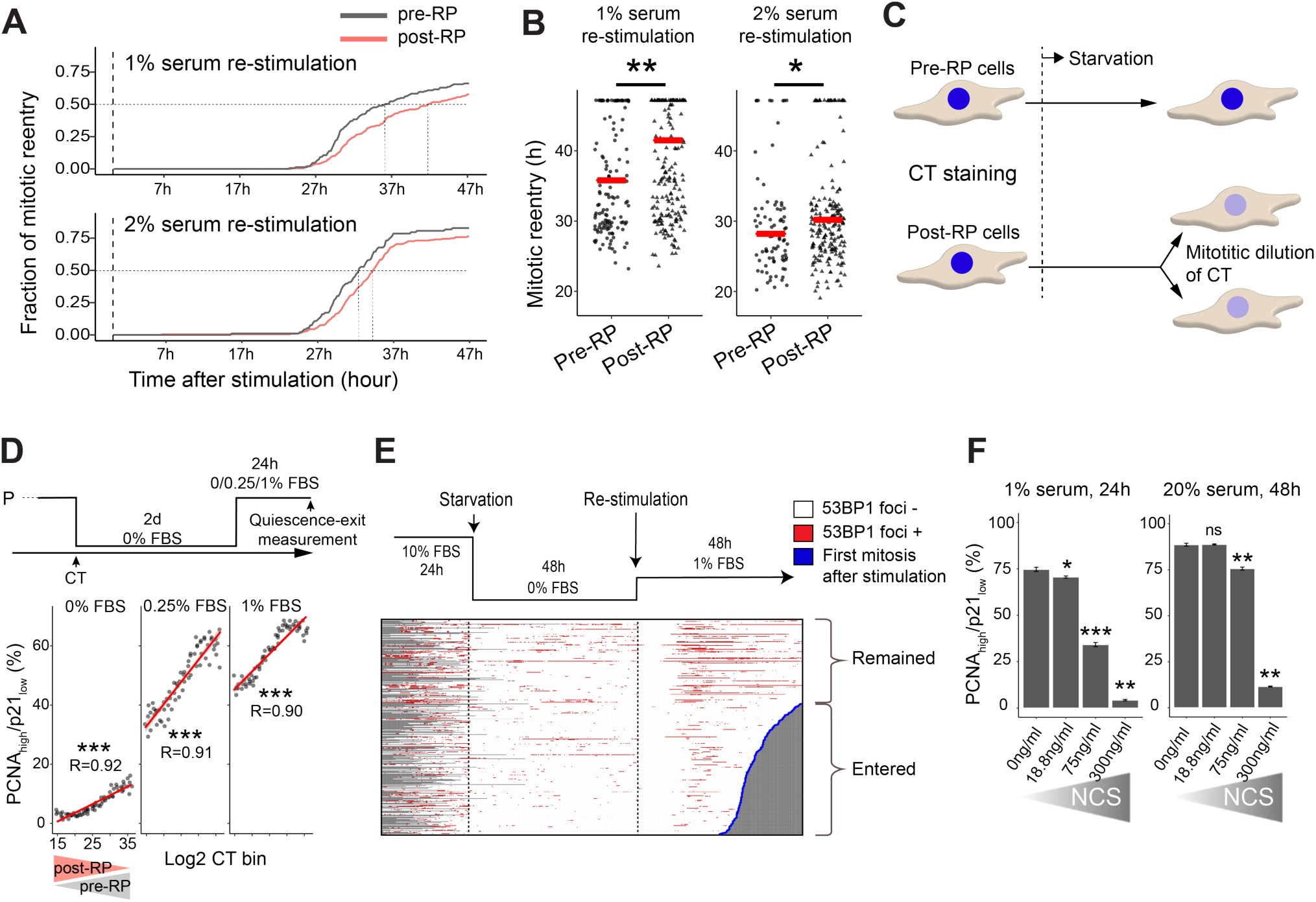
DSBs drive cells into deeper quiescence. (A, **B)** Fraction (A) and timing (B) of mitotic reentry in growth re-stimulated quiescent cells of pre-RP and post-RP origins, analyzed from Fig. 2G and Fig. S2F. In (B), * and ** indicate p < 0.05 and 0.01 in Mann Whitney U test. **(C)** Experimental scheme for distinguishing pre-RP and post-RP cells using CT dye. **(D)** Quiescence depth measurement in pre-RP and post-RP origin cells in the CT dye-based experiment. Top: experimental design. Cells were binned according to their CT intensity, and the percentage of PCNA_high_/p21_low_ cells was calculated per CT bin (n = 3). Red lines show linear fits. ** and *** denote p < 0.001 and p < 0.0001 by Spearman’s rank correlation. **(E)** 53BP1 foci dynamics in cells that remained quiescent or reentered the cell cycle – data from Fig. 2G (n > 380 cells). Cells were aligned by mitotic timing (indicated by the blue line) after the growth re-stimulation. Color scheme as in Fig. 2G. **(F)** Quiescence depth measurement in cells treated with NCS. Two-day starved quiescent cells were treated with the indicated NCS concentrations for 1 hour, then growth stimulated (n = 2). Percentage of PCNA_high_/p21_low_ cells for each condition is shown. *, **, and *** indicate p < 0.05, < 0.01, and < 0.001 by one-tailed *t*-test.

As an orthogonal test, we used a CellTrace (CT) Violet dye-labelling method^38^ to separate post-RP and pre-RP cells. After the initial labeling, the dye is diluted by cell division, and thus post-RP cells (which divide before entering quiescence) show low CT intensity, whereas pre-RP cells (which do not divide) retain high CT intensity (Fig. 3C, Fig. S3A). After growth restimulation, quiescent cells with low CT intensity (enriched for post-RP origin) displayed a lower tendency to reenter the cell cycle (becoming PCNA_high_/p21_low_) than high CT intensity cells (enriched for pre-RP origin) (Fig. 3D, 0.25% and 1% FBS). Consistently, a minor perturbation of refreshing the starvation medium (0% FBS) induced a small fraction of high CT intensity (pre-RP origin) cells, but not low CT intensity (post-RP origin) cells, to exit quiescence (Fig. 3D). These results indicate that quiescent RPE cells of post-RP origin reside in a deeper quiescent state than those of pre-RP origin.

Next, we examined the relationship between DSBs and cell division after growth restimulation. Cells that failed to divide (deep quiescence) exhibited more 53BP1 foci during restimulation compared to those that divided (shallow quiescence) (Remained vs Entered, Fig. 3E, and Fig. S3B). Furthermore, direct induction of DSBs in quiescent cells via NCS treatment resulted in a dose-dependent increase in quiescence depth, as evidenced by a decreased fraction of PCNA_high_/p21_low_ cells after restimulation (Fig. 3F). Taken together, these results suggest that DSB accumulation—particularly in cells that pass the RP at the onset of quiescence induction—deepens quiescence and impedes cell cycle reentry and division upon growth restimulation.

### DNA-PK promotes, while ATM attenuates, cell cycle reentry of DSB-containing quiescent cells

In line with previous findings that DNA damage is repaired upon quiescence exit in HSCs^17^, we observed that DSBs in quiescent cells were gradually repaired after growth restimulation (Fig. 4A). Nevertheless, this repair is not sufficient to completely prevent the negative effects of DSBs on cell cycle reentry, as DSBs still impede this process (Fig. 3). These observations prompted us to examine how the multiple components of the DDR—including DNA repair and checkpoint response—contribute to the reentry process.

**Figure 4:**
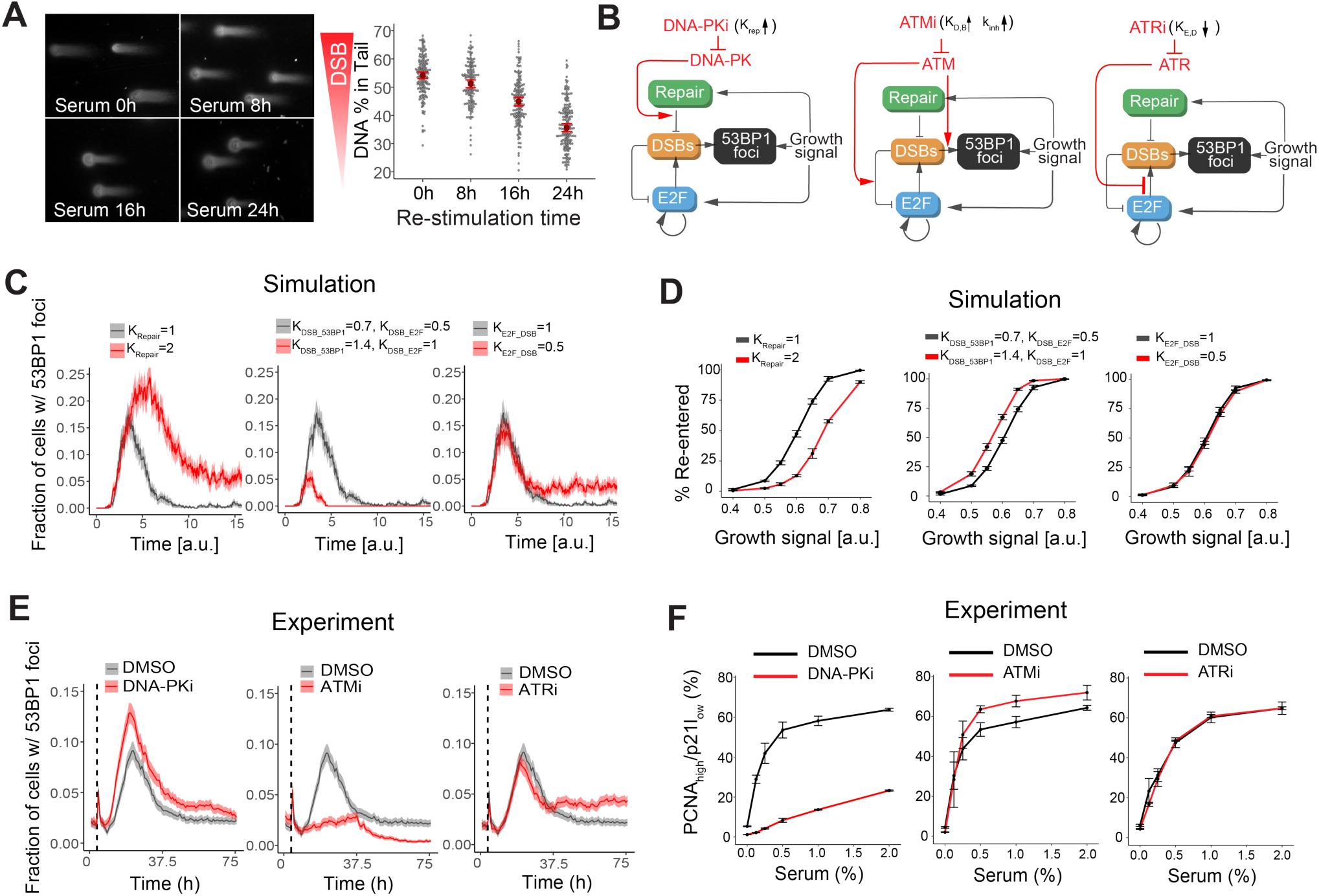
DNA-PK promotes, while ATM attenuates, quiescence exit. **(A)** Representative neutral comet images of cells exiting quiescence (n > 100 cells per condition). Percent DNA in the comet tail is also shown. Two-day starved cells were growth-stimulated with 4% FBS for up to 24 hours and analyzed by neutral comet assay. **(B)** Diagram of perturbation targets for each PIKK inhibitor. Known functions of each PIKK and corresponding parametric changes used to mimic inhibition in the model are indicated. **(C)** Simulated effects of PIKK inhibition on the fraction of cells with 53BP1 foci after re-stimulation of quiescent cells. 53BP1 activity above 0.8 was considered positive for foci (threshold set to match experimental results in Fig. 2H). Black and red lines indicate non-inhibited and inhibited scenarios, respectively. Results from 500 stochastic simulations. Mean and 95% confidence interval from bootstrapping are shown in thick lines and colored ribbons, respectively. **(D)** Simulated percentage of cells re-entering the cell cycle under PIKK inhibition (n = 3; 500 simulated cells per condition). E2F > 0.1 at model time 10 was considered as cell cycle re-entry. Black and red indicate non-inhibited and inhibited scenarios, respectively. Error bar, SEM. **(E)** 53BP1 foci dynamics in growth-stimulated RPE-53BP1 cells treated with PIKK inhibitors (n > 3500 cells per condition per time point). Two-day starved RPE-53BP1 cells were stimulated with 1% FBS and treated with ATMi (1μM Ku-60019), ATRi (1μM VE-821), or DNA-PKi (2μM NU-7441). Imaging was performed every 45 minutes, and the fraction of cells with 53BP1 foci was calculated at each time point. Vertical dashed line marks re-stimulation. Mean and 95% confidence interval from bootstrapping are shown in thick lines and colored ribbons. **(F)** Percent of PCNA_high_/p21_low_ cells after 24 hours of growth re-stimulation with PIKK inhibition (n = 3). Two-day starved cells were re-stimulated at the indicated serum concentration. Inhibitor concentrations as in (E). Error bar, SEM.

**Figure 5:**
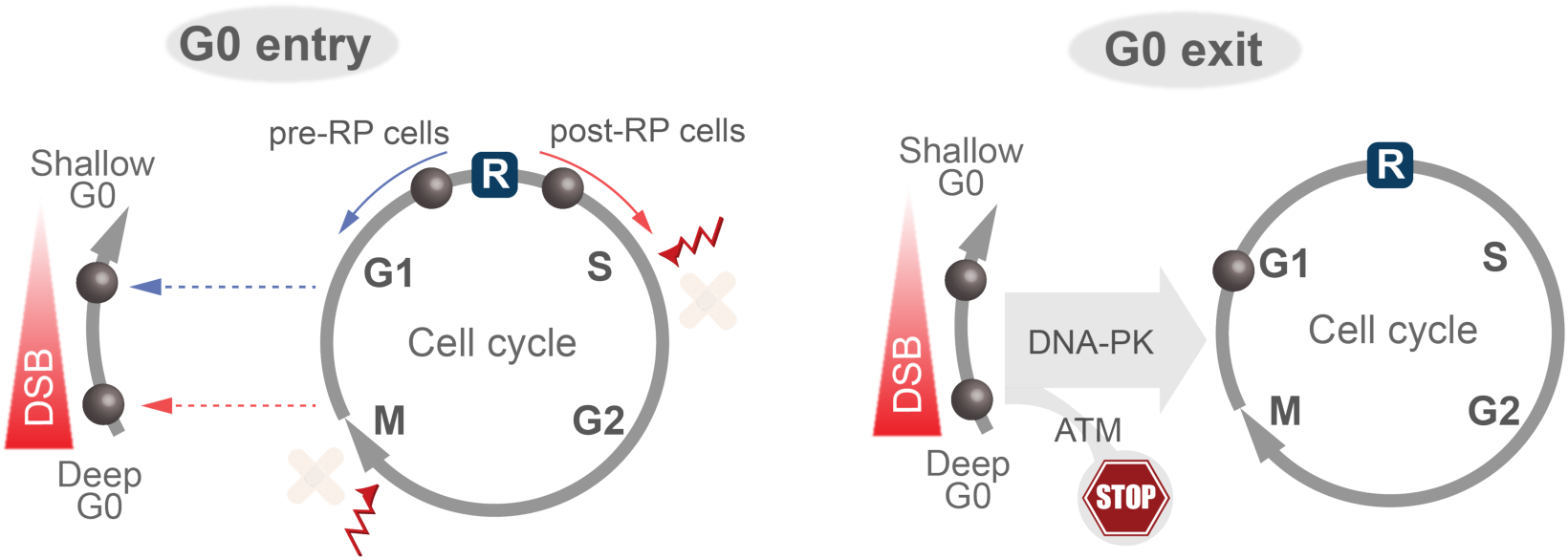
The working model. Upon quiescence induction by starvation, post-RP cells undergo cell cycle activity with limited DNA repair and accumulate DSBs, resulting in deep quiescence. In contrast, pre-RP cells withdraw from the cell cycle and avoid DSB accumulation, resulting in shallow quiescence. Accumulated DSBs activate DDR components that affect quiescence exit: DNA-PK facilitates DSB repair and promote quiescence exit, while ATM delays or inhibits quiescence exit when DSB levels are high.

To dissect the role of DDR during quiescence exit and cell cycle reentry, we first perturbed the function of key PI3K-like kinases— DNA-PK, ATM, and ATR—in our computer model. DNA-PK is essential for NHEJ^13^, the canonical DSB repair pathway in G0 and G1 cells. ATM activates DNA damage checkpoints^13^ and is required for 53BP1 foci formation^39^. ATR is not directly activated by DSBs but prevents their formation by limiting replication fork collapse^13^. We emulated the inhibition of these kinases *in silico* by adjusting the corresponding parameter values (Fig. 4B). Simulation results suggested that DNA-PK and ATM have opposing roles during quiescence exit. Inhibiting DNA-PK reduced DSB repair, amplified the 53BP1 pulse (Fig. 4C, left; Fig. S4), and hindered quiescence exit (Fig. 4D, left; Fig. S4). By contrast, inhibiting ATM, by disrupting 53BP1 foci formation^39^ and DNA-damage checkpoint^13^, reduced 53BP1 pulse (Fig. 4C, middle; Fig. S4) and facilitated quiescence exit (Fig. 4D, middle, Fig. S4). ATR inhibition appeared to have little effect on 53BP1 pulse (Fig. 4C, right; Fig. S4) and quiescence exit (Fig. 4D, right; Fig. S4), consistent with its effect on DSBs and 53BP1 only after cell cycle reentry and E2F activation.

To experimentally test these model predictions, we treated starvation-induced quiescent RPE cells with growth restimulation under specific PIKK inhibitors, and monitored 53BP1 foci formation using time-lapse imaging (Fig. 4E) as well as quiescence exit using end-point FACS analysis (Fig. 4F). Our experimental observations were in good agreement with model predictions in the overall trend: inhibiting DNA-PK amplified the 53BP1 pulse (Fig. 4E, left) and hindered quiescence exit (Fig. 4F, left), whereas inhibiting ATM suppressed the 53BP1 pulse (Fig. 4E, middle) and facilitated quiescence exit (Fig. 4F, middle), and inhibiting ATR had little effects on the 53BP1 pulse (Fig. 4E, right) and quiescence exit (Fig. 4F, right).

## DISCUSSION

Our study demonstrates that quiescence induction rapidly triggers the accumulation of DNA double-strand breaks in a cell cycle stage-dependent manner, with post-restriction point cells accumulating more DSBs and consequently entering deeper quiescence states. Numerous studies on adult stem cells have found that long-term wear and tear during aging contribute to the accumulated DNA damage in quiescent cells^17,23,40^. The present study demonstrates that during quiescence entry, accumulation of DNA damage (DSBs) can occur rapidly, within hours of quiescence induction.

Not only is the rapid accumulation of DNA damage in quiescent cells (hours vs. months or years) surprising, but the order of action is too: DNA damage has been considered the cause, rather than the consequence, of cell cycle arrest—including “spontaneous” quiescence induced by endogenous DNA damage under non-perturbed conditions^9,10,41^. Now we show the previously unrecognized reverse relationship: quiescence induction itself causes the accumulation of DNA damage. The fact that starvation alone, without continued cell cycle activity, did not lead to the accumulation of DSBs (Fig. 2C), and that cells under contact inhibition without starvation also exhibited DSB accumulation during quiescence entry (Fig. S1C, D), indicates that DSB accumulation is not merely a response to starvation. Rather, it is intrinsically associated with the process of cell cycle exit and quiescence entry.

Mechanistically, our experiments and modeling suggest that DSB accumulation following quiescence induction is not due to increased DSB generation, but is primarily driven by continued DNA damage-prone cell cycle activity coupled with reduced DNA repair under growth restriction conditions during quiescence entry. Consequently, DSBs accumulate predominantly in post-RP cells at the time of quiescence induction (Fig. 2G, H). However, other sources of DNA damage should also contribute, as pre-RP cells are not entirely DSB-free. These might include metabolic shifts^42^ to oxidative phosphorylation, which can generate ROS, or large-scale transcriptional reprogramming^43^ that could introduce topological stress on DNA, which, if not resolved, leads to DSBs.

DSB accumulation triggered by quiescence induction links cell cycle position at the time of quiescence induction to the observed variability in quiescence depth. Previously, we found in rat embryonic fibroblast (REF) cells that the depth of quiescent cells reflected a memory of their preceding cell cycle position relative to the RP, though the underlying mechanism remained unclear^38^. Our current findings suggest that the differential DSB accumulation during quiescence entry likely contributes to this memory effect of prior cell cycle position.

Our study suggests a critical—yet counterintuitively genome-destabilizing—role for DSB repair during quiescence exit and cell cycle reentry. Our data suggests that DSB repair facilitates quiescence exit (Fig. 4F) by removing the triggers for ATM-enforced DNA damage checkpoint activation. However, this DNA-PK-mediated repair is less faithful than HR, which is unavailable during quiescent exit due to the absence of a sister chromatid template, and may introduce small indels at the repair sites through NHEJ or related pathways^44^. Together with findings that quiescence exit can provoke *de novo* DNA damage^45^ and origin underlicensing^46^, these observations highlight the challenges and risks of mutagenesis and genome instability that accompany the transition from quiescence to proliferation.

Key molecular details underlying DSB and cell cycle regulation during quiescence induction and exit remain unknown. First, how cells down-regulate DSB repair under growth restriction conditions is unclear. Although E2F activity does not turn off while cells complete the current cell cycle—reflecting the bistable and hysteretic properties of E2F expression^47,48^—it may fall below the threshold required to sustain the expression of E2F-targeted DSB repair genes. This could provide an explanation but needs to be validated. Second, the components beyond ATM that underlie the DSB checkpoint during quiescence exit remain elusive. Since ATM activates p53, the checkpoint could be mediated by a p53-dependent mechanism^10^.

One important question that emerges from our work is how DSB accumulation associated with quiescence induction impacts cellular aging *in vivo*. DSBs accumulated during quiescence entry drive deep quiescence, serving as a direct entry point into a pre-senescent state^35^, whereas repeated cycles of quiescence entry and exit could gradually promote cellular aging through the accumulation of mutations caused by recurring DSBs and error-prone repair. It is possible that circadian-orchestrated quiescence exit/entry, observed in several cell types^49–53^, may mitigate such genome deterioration and quiescence deepening by synchronously positioning cells in the pre-RP state at quiescence entry. However, whether pre-RP cells indeed predominate during such synchronized quiescence entry in tissues is yet to be studied. Further investigation of DSB accumulation and repair in quiescent cells will be essential for understanding how tissue homeostasis and aging are shaped in our body.

## LIMITATION

This study was based on a widely used non-cancerous human cell line, RPE-1, and the generalization of our findings needs to be tested in other cell contexts in future studies. The rates of endogenous DSB generation and repair were not separately measured, as live-cell reporters for these two processes are currently lacking.

## RESOURCE AVAILABILITY

Lead contact: Requests for further information and resources should be directed to and will be fulfilled by the lead contact, Guang Yao

Materials availability: All unique/stable reagents generated in this study are available from the lead contact with a completed materials transfer agreement.

Data and code availability: All original code has been deposited at [https://github.com/KotaroFuji/GFDamageSim] and will be publicly available as of the date of publication. Any additional information required to reanalyze the data reported in this paper is available from the lead contact upon request.

## ACKNOWLEDGEMENT

We thank Alexis Barr for kindly providing us with the clonal RPE-PCNA-p21 cell line and its parental RPE-hTERT cell line. K.F. thanks Kathleen Anne Lasick and Julie Huynh for helping

K.F. troubleshoot occasional microscopy hiccups. We thank Kimiko Della Croce for technical suggestions for the comet assay. We thank Andrew Capaldi, Nathan Ellis, and Ted Weinert for constructive comments throughout the project. This research was partially supported by National Science Foundation Grants DMS-1463137 and DBI-2016035 (to G.Y.).

## AUTHOR CONTRIBUTION

Conceptualization, Methodology, Investigation, Formal Analysis, Visualization, Writing – Original Draft, Review & Editing, K.F; Supervision, Writing – Review & Editing, A.P; Project Oversight, Supervision, Writing – Review & Editing, Funding Acquisition, G.Y.

## METHODS

### Cell lines

The clonal retinal pigment epithelial cell line with the PCNA and p21 reporters^10^ used in this study (RPE cells for short) was a kind gift from Dr. Alexis Barr. RPE cells with the 53BP1 reporter^30^ (RPE-53BP1 cells for short) are from a single-cell clone derived from RPE-hTERT parental cells (a gift from Dr. Alexis Barr) stably integrated the plasmids Apple-53BP1trunc (Addgene, #69531) and pLentiPGK-DEST-H2B-iRFP670 (Addgene, #90237).

### Cell Culture and quiescence induction

Cells were passaged every 2-3 days and maintained at subconfluency in growth medium: Dulbecco’s Modified Eagle’s Medium (DMEM with sodium pyruvate additive) supplemented with 10% fetal bovine serum (FBS; SIGMA, F0926) and 4 mM GlutaMax (Gibco, 35050-061). To induce quiescence, asynchronously growing cells at ∼50% confluence on 6-well or 12-well plates in growth medium were washed twice with DMEM and cultured in DMEM for 2 days. In microscopy experiments, FluloroBrite DMEM (Gibco, A1896701; added with 1 mM sodium pyruvate) and 12-well/35 mm glass-bottom plates/dishes (Fisher Scientific, NC0799106; MatTek, P35GCOL-1.5-14-C) were used for the culture.

### Neutral comet Assay

The following protocol was adapted and revised from Trevigen’s instruction for the CometAssay kit (Trevigen, 4250-050-K) to accommodate the differences in the micro-well slides, buffers, and an electrophoresis tank. Briefly, harvested cells were washed once with and resuspended in ice-cold DPBS, and the cell suspension was quickly mixed with pre-warmed and molten light-melting agarose. The cell-LMA suspension (1.25% LMA) was applied on micro-well slides (Electron Microscopy Sciences, 63423-08) precoated with normal-melting agarose, and a covered slide (VWR, 48368-062) was put on top to allow the gel suspension to form a uniform layer. The slides were put in 4° C until the gelation, and they were lysed in lysis solution (2.5 M NaCl, 0.1 M EDTA, 10 mM Tris, 1% w/v Lauryl Sarcosine, 1% v/v Triton X-100) at 4 °C overnight. Subsequently, slides were immersed in neutral electrophoresis buffer (100 mM Tris, 496 mM Sodium Acetate, pH adjusted to 9.0 by glacial acetic acid) for 30 minutes without a current and for 40 minutes with a current (1.15 V/cm) in a dark 4 °C cold room. Slides were immersed in DNA precipitation solution (1 M NH_4_Ac in 85% EtOH) and 70% EtOH for 35 minutes each at room temperature. Slides were completely dried at 37°C and stained with SYBR Gold (Invitrogen, S11494) for 30 minutes, after which they were briefly washed with water, dried, and viewed under a microscope to quantify the comet tails. Comet signals were quantified automatically using OpenComet^54^ in Fiji, an extended version of Image J.

### Time-lapse imaging

Cells were imaged on a Nikon Eclipse Ti-E microscope fitted with a Hamamatsu Orca-Flash 4.0 V2 camera. Samples were enclosed in an OKO labs incubation chamber to maintain humidity, 37 °C temperature, and 5% CO2.

### Quiescence depth assay

To assess quiescence depth, 2-day starved cells were switched from starvation medium (DMEM without FBS and L-glutamine) to growth-stimulation medium (DMEM containing 4 mM GlutaMax and the indicated concentration of FBS, except noted otherwise) and cultured for the indicated duration. Cells were subsequently trypsinized and fixed with 1% paraformaldehyde for flow cytometry. Cells that exited quiescence and reentered the cell cycle were determined based on their PCNA_high_/p21_low_ profiles, as shown in Fig. S1A. Quiescence depth is determined by the serum threshold or duration required to activate cells to reenter the cell cycle^33–35^. At a given serum concentration, a smaller percentage of deeper quiescent cells are able to reenter the cell cycle by a given time than that of shallower quiescent cells. See text for details.

### CellTrace Violet staining and cell-cycle phase analysis

For CellTrace Violet (Invitrogen, C34557) staining, cells were processed according to the manufacturer’s protocol. Briefly, cells were stained with 6 μM CellTrace Violet in pre-warmed DPBS and incubated for 20 minutes at 37°C. Cells were washed twice with 10% FBS DMEM before being put into the desired medium. For cell-cycle phase analysis, cells were pulse-labeled with 10 μM EdU for 20 minutes before the harvest. Harvested cells were fixed with 4% paraformaldehyde for 15 minutes at RT, after which they were permeabilized by washing-permeabilization buffer (1% BSA and 0.1% Tween-20 in DPBS) and processed for Click-iT reaction. Cells were washed once with washing-permeabilization buffer before being put into DNA-staining solution (250 μg/ml RNase, 1 μg/ml Hoechst 33342, and 1% BSA in DPBS).

### qPCR

Total RNA was extracted using Quick-RNA Mini Prep (ZYMO research, R1054) according to the manufacturer’s protocol. The quality of extracted RNA was confirmed using NanoDrop. cDNA was synthesized using Maxima First Strand cDNA Synthesis Kit for RT-qPCR, with dsDNase (Thermo Scientific, K1672) according to the manufacturer’s protocol. qPCR was conducted in ABI 7300 using Power SYBR Green PCR Master Mix (Applied Biosystems, 4367659).

### Mathematical Modeling

The purpose of our model was not to simulate detailed molecular interactions, but rather to capture the dynamics of DSBs and key cellular activities in the context of growth signaling, with the aim of clarifying how DSBs and quiescence entry and exit influence each other.

We developed a minimal model incorporating four key entities—DSBs, DNA repair machinery, E2F, and 53BP1—and their interactions with growth signaling. Growth factor (GF) activates E2F^47^, a transcription factor crucial for proliferation^55^ that is regulated by positive feedback loops, conferring ultrasensitivity^47,48^. E2F stimulates proliferative activities that increase DSBs^1–5^. These DSBs are repaired by DNA repair machinery in a GF-dependent manner^15,16,20^. Unrepaired DSBs inhibit E2F activity^10^. In our model, 53BP1 reflects both DSB levels and GF concentration.

These relationships were implemented as delay differential equations (DDEs) below.

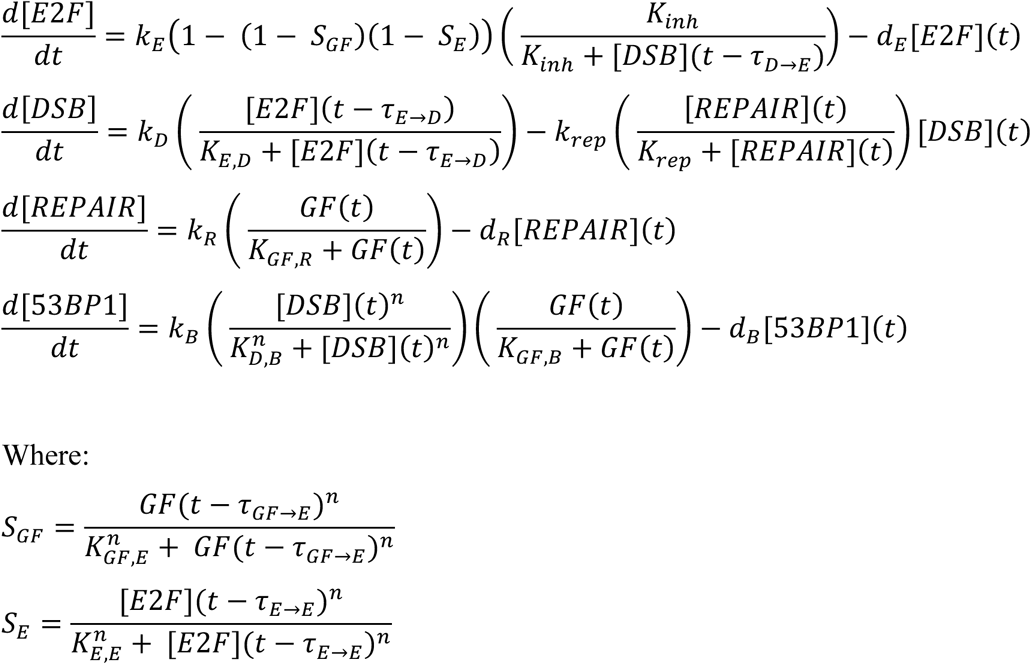

Time delays, denoted by τ, were implemented to account for different regulatory time scales (Table S1). For example, GF to E2F activation requires many hours, as it involves transcription and translation of multiple factors in multiple steps, whereas DNA repair can be completed within an hour. To confer ultrasensitivity to E2F and 53BP1 activation (activation of 53BP1 in the model is equivalent to 53BP1’s recruitment to DSB site), we set a hill coefficient for relevant parts. GF was altered in a stepwise fashion to mimic starvation and restimulation.

Simulations were performed in Python with parameters specified in Table S1. When necessary, we implemented stochasticity using the chemical Langevin approach^56^, as in our previous Rb-E2F network model^34^.

To evaluate the robustness of 53BP1 pulsing, we varied the Michaelis-Menten constant, synthesis rate, and degradation rate of all four components—doubling or halving each parameter and simulating all 16,384 (2^14^) possible combinations. Across these conditions, pulsatile 53BP1 dynamics following growth stimulation were a typical outcome (Fig. S2E).

### Image analysis

The cells were semi-automatically tracked using custom MATLAB scripts used in a previous study^57^. 53BP1 foci were quantified using custom python scripts. Briefly, cellular nuclei were segmented using otsu method implemented in the python scikit-image package^58^. Denoised and gamma-corrected 53BP1 reporter signal was used to determine nuclear regions (i.e., foreground) from the rest (i.e., background). 53BP1 foci were detected using the blob_log feature in the sckit-image package^58^. To analyze the effect of the prior cell cycle positions on subsequent DSBs levels, we limited our analyses to actively cycling cells at the time point of starvation. Thus, pre-arrested cells, such as senescent-like (i.e., irreversibly arrested cells) and quiescent-like cells^9,10,59^ (i.e., cells with reversible G0 arrest), at the time point of starvation were removed from the tracked results. See the section below for more details.

### Filtering out pre-arrested cells from time-lapse imaging data

To assess how prior cell cycle status affects DSB levels after starvation, we analyzed only actively cycling cells at the onset of starvation, excluding spontaneously quiescent cells (which are present under normal conditions due to endogenous DNA damage^9,10,59^). First, cells without any mitosis during the 5-day experiment were removed. Next, we excluded cells that divided before but not after starvation and whose time from mitosis to starvation exceeded the typical time from mitosis to the restriction point (RP), since such cells would have been expected to commit to another division unless already pre-arrested. We estimated this critical interval (M-RP_time_) by time-lapse imaging PCNA-mRuby/p21-GFP RPE cells, tracking over 300 cells during the transition from growth to starvation. From G1 duration and the ratio of G1 cells that divided vs. did not divide post-starvation, we determined M-RP_time_ to be 8.4 hours. Thus, G1 cells with mitosis-to-starvation time > 8.4 hours and no mitosis post-starvation were excluded. These filtered cells showed pre-existing DNA damage (53BP1 foci), validating our approach (Fig. S5).

## SUPPLEMENTAL INFORMATION

Document S1: Figure S1-S5, Table S1

---

**Figure S1:**
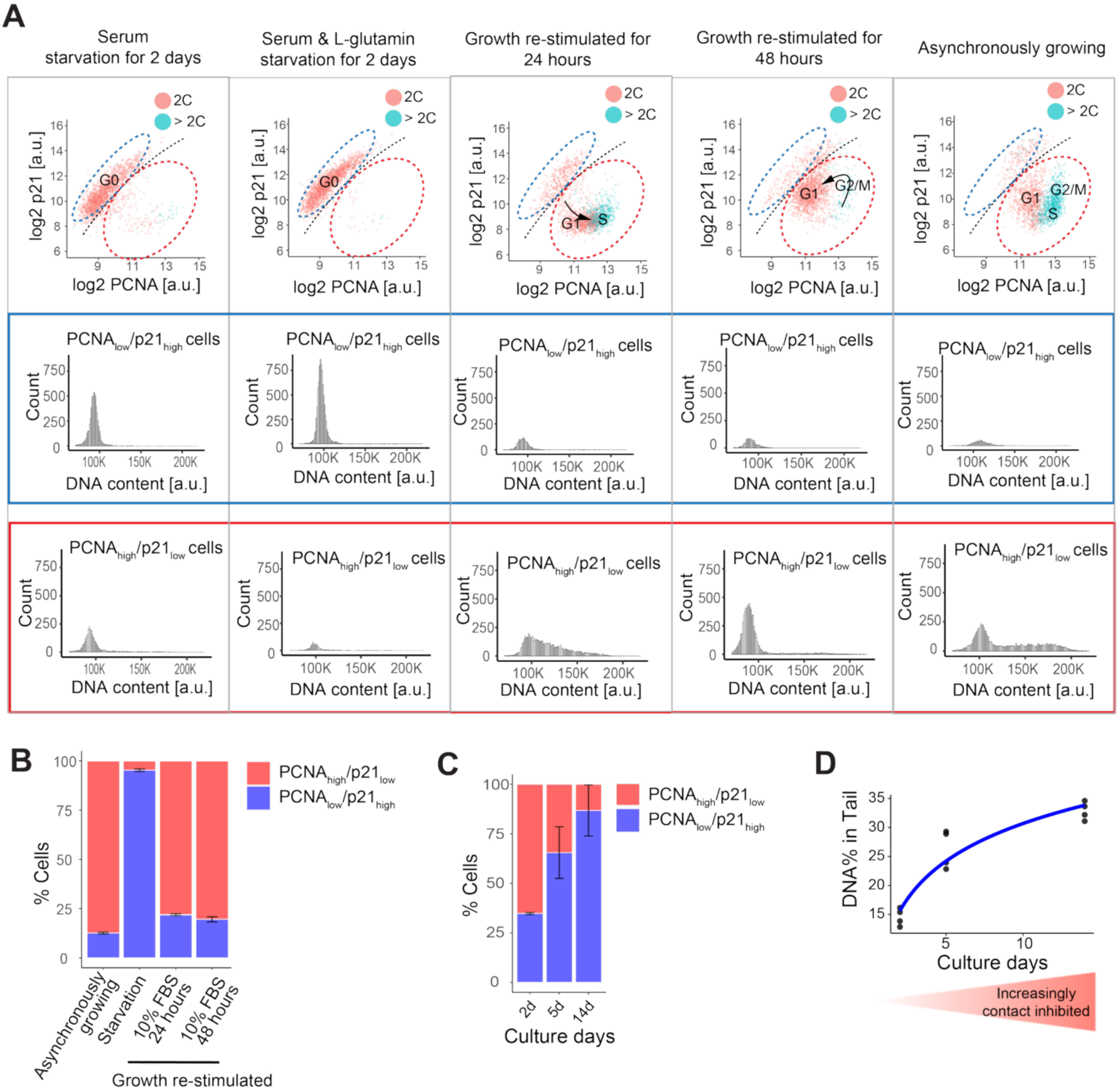
Quiescence induction evokes DSBs, Related to. Figure 1**. (A)** Top: PCNA and p21 profiles of RPE cells under the indicated culture conditions. Growth-stimulation (10% FBS) was applied to cells after 2 days of serum & L-glutamine starvation. Red and green dots indicate cells with 2C and >2C DNA content, respectively. Blue and red dashed lines demarcate PCNA_low_/p21_high_ and PCNA_high_/p21_low_ populations, respectively. Histograms show DNA-content profiles for PCNA_low_/p21_high_ cells (blue box, middle) and PCNA_high_/p21_low_ cells (red box, bottom) under each culture condition. **(B)** The percentage of PCNA_low_/p21_high_ cells and PCNA_high_/p21_low_ cells in asynchronously cycling, starved, or growth-stimulated cells (n = 3). **(C)** The percentage of PCNA_low_/p21_high_ cells and PCNA_high_/p21_low_ cells in cells cultured under increasingly confluent conditions (n = 3). **(D)** The percent of DNA in the tail, based on neutral comet assay, in cells cultured under increasingly confluent conditions (n=4).

**Figure S2:**
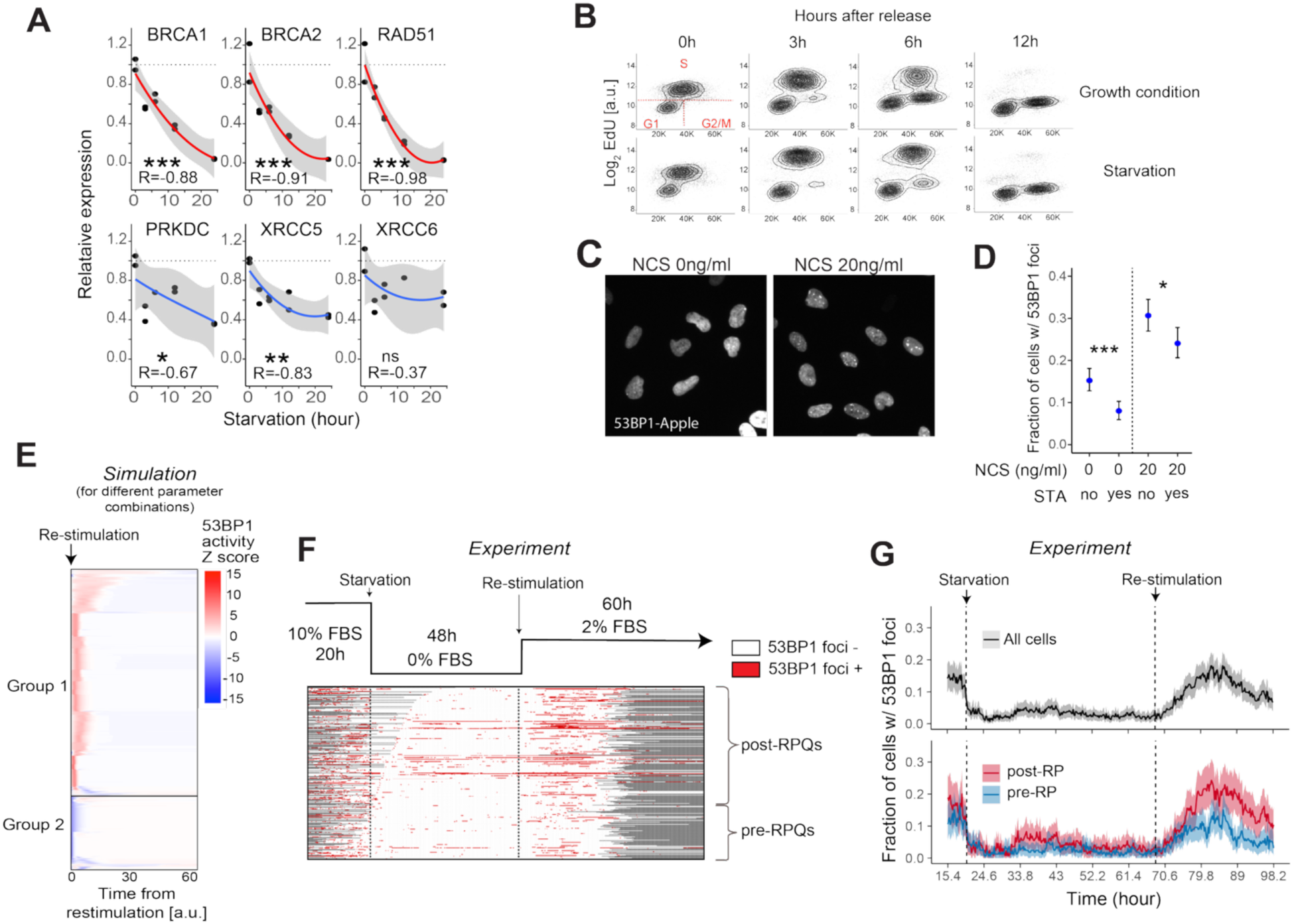
Growth-dependent DDR and 53BP1 dynamics in cells entering and exiting quiescence, Related to. Figure 2**. (A)** Gene expression of HR and c-NHEJ genes in cells starved up to 24 hours (n =2). BRCA1, BRCA2, and RAD51 belong to the HR pathway and PRKDC, XRCC5, and XRCC6 to c-NHEJ. *, **, and ***, p < 0.05, 0.01, and 0.001, respectively (Spearman’s correlation). **(B)** Cell-cycle-phase analysis on cells released from the HU arrest in starvation or growth condition. Cells were first arrested by HU in a growth condition (10% FBS) for 24 hours and released from the arrest in starvation or growth condition for up to 12 hours. **(C)** Representative images of cycling RPE-53BP1 cells treated with 20ng/ml NCS for an hour. **(D)** The fraction of cells with 53BP1 foci in RPE-53BP1 cells treated with NCS with or without starvation (n > 500 cells per condition). Starvation was applied 4 hours before the imaging. NCS was directly added to the medium at the indicated concentration 1 hour before the imaging. Error bar, 95% confidence interval from bootstrapping. * and ***, p-value < 0.05 and 0.001, respectively (Fisher’s exact test). **(E)** 53BP1 dynamics following growth re-stimulation under varied parameter conditions. Michaelis-Menten parameters, synthesis rates, and degradation rates for all four nodes were systematically doubled or halved from default values and combined (2^14^ = 16384 combinations; see Table S1 for parameters). For each parameter set, 53BP1 activity dynamics after growth re-stimulation were simulated using the deterministic model. Color indicates 53BP1 activity Z-score per row (53BP1 activity normalized across time). Simulations with pulsatile 53BP1 activity were classified as Group 1, while those with non-pulsatile 53BP1 activity were classified as Group 2. **(F)** 53BP1 foci dynamics in RPE-53BP1 cells entering and exiting quiescence (n > 310 cells). **(G)** Fraction of cells with 53BP1 foci over time in pre- and post-RP groups – data from G (n > 90 and 220 cells for pre- and post-RP, respectively). Median and 95% confidence interval from bootstrapping are respectively shown in thick lines and colored ribbons. Only time frames where more than 80% of all tracked cells have valid data are included.

**Figure S3:**
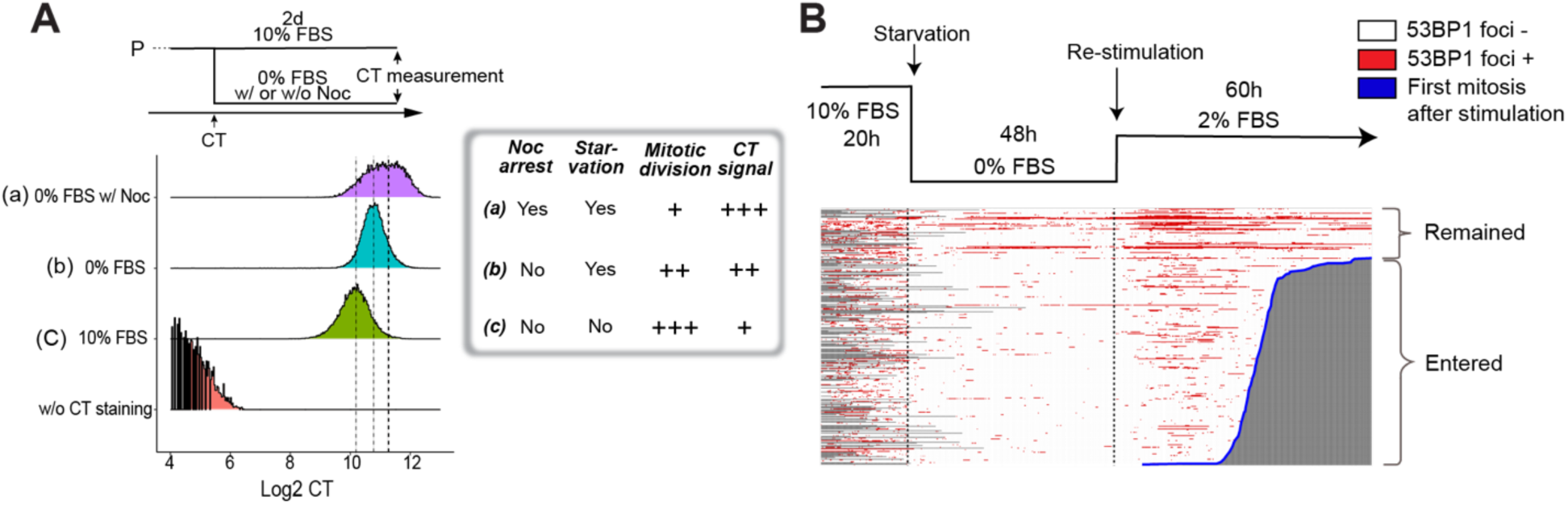
Higher DSB levels correlate with deeper quiescence, Related to. Figure 3**. (A) CellTrace (**CT) intensities under conditions with different mitotic propensities. The experimental design is depicted at the top. Cycling cells were stained with CT, washed off of CT, and subsequently cultured in (a) starvation condition with 120nM nocodazole (0% FBS w/ Noc), (b) starvation condition (0% FBS), or (c) growth condition (10% FBS). Noc: nocodazole. The inset table summarizes the culture treatment, mitotic propensity, and CT signal in each of the three conditions. **(B)** The 53BP1 foci dynamics in cells remained quiescent or reentered the cell cycle under 2% FBS stimulation – data from Fig. S2F (n > 310 cells). Cells were aligned according to their mitotic timing after the growth stimulation, as in Fig. 3E.

**Figure S4:**
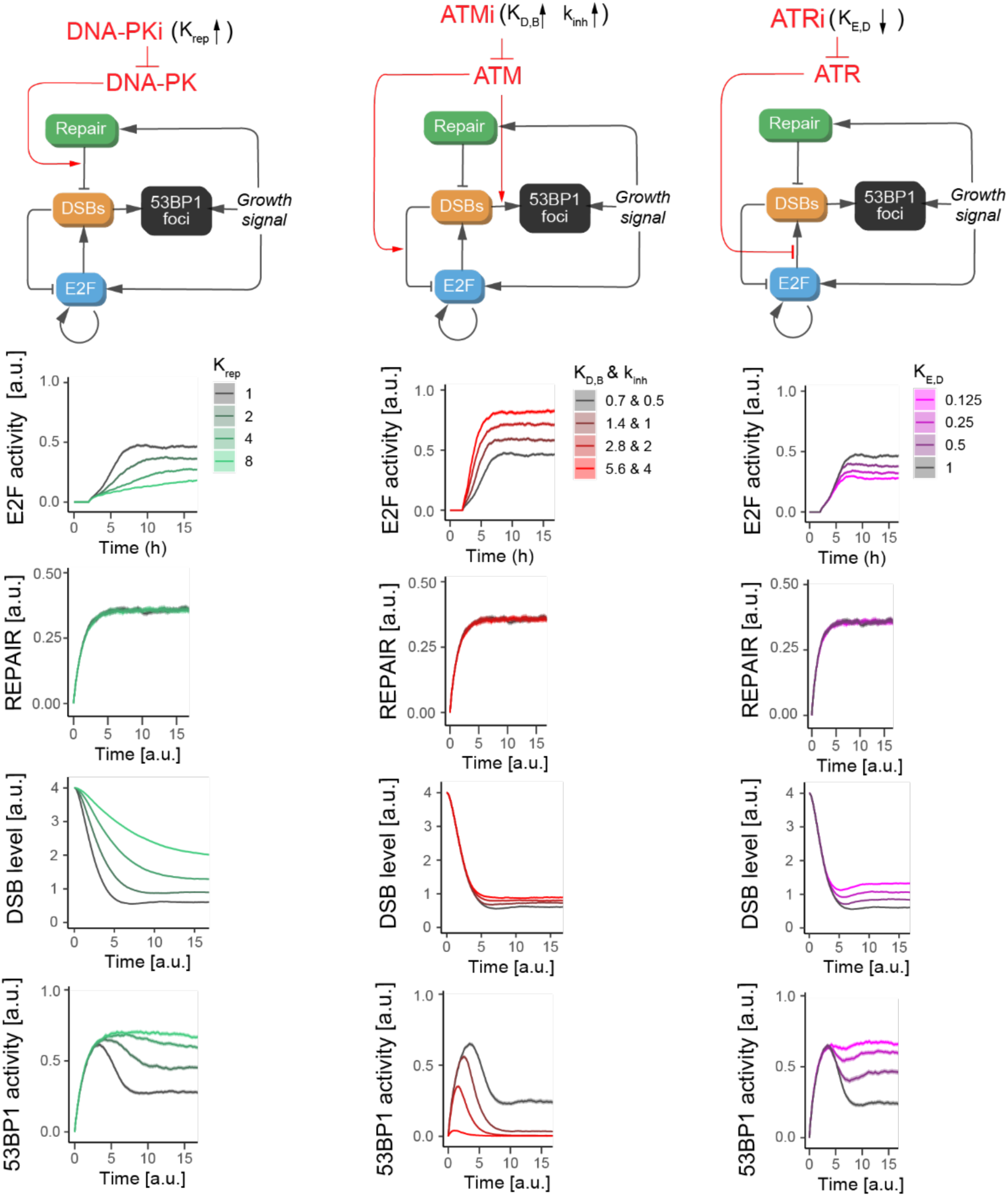
The role of DDR in quiescence exit, Related to. Figure 4. The model was simulated under different perturbation conditions that mimic PIKK inhibitions, and the dynamics of the model components were analyzed. Each parameter was altered in the indicated direction from the default value (τ : increase, τ: decrease) in log_2_ intervals.

**Figure S5:**
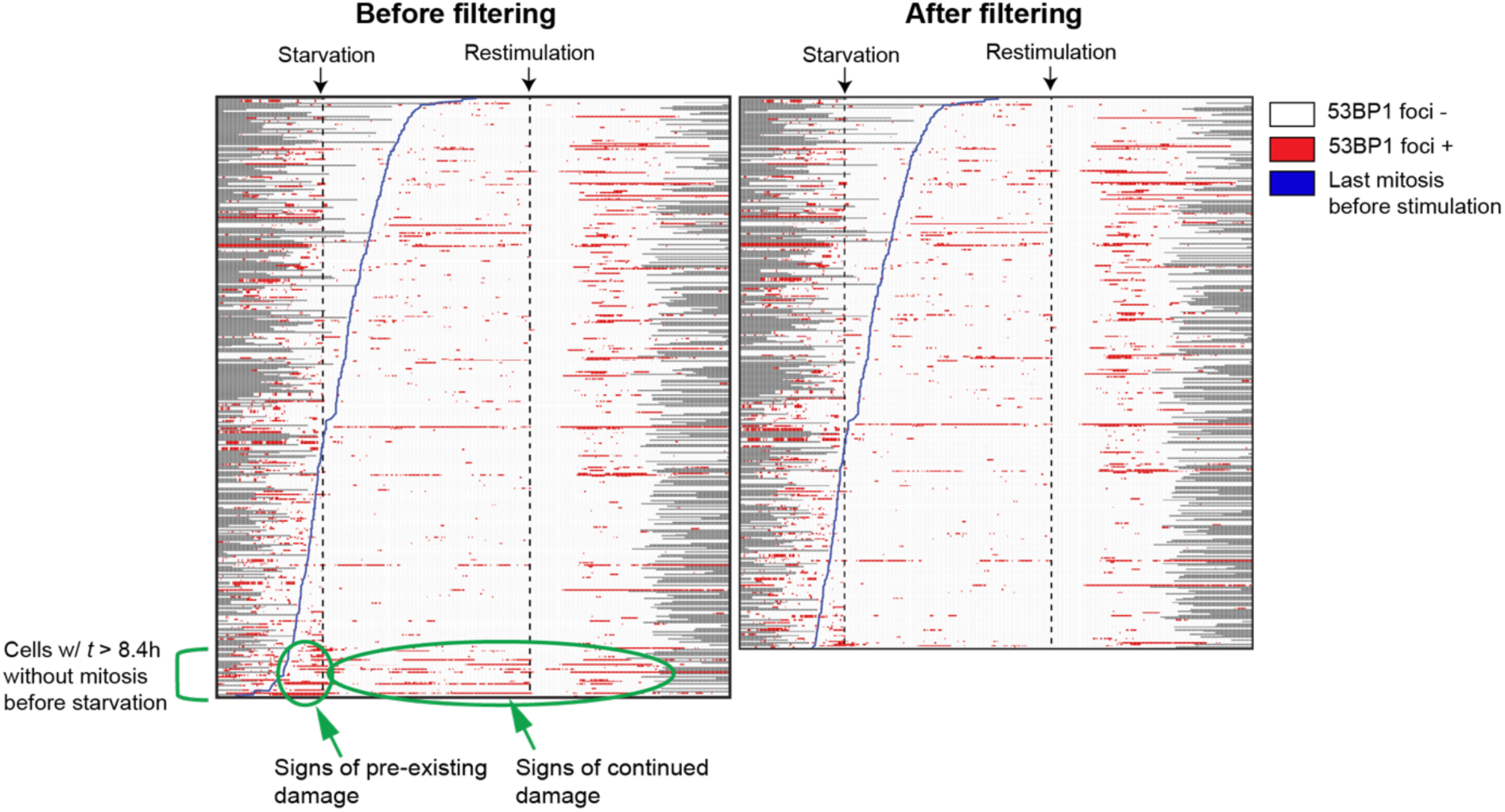
53BP1 profile of filtered-out cells, Related to Methods. Cells that spent longer than 8.4 hours without mitosis before starvation and did not divide after starvation were considered pre-arrested and excluded from analysis (see *Methods* for details).

**Table S1:** Model parameters, Related to Methods.

| Symbol | Values | Description |
| --- | --- | --- |
| $k_E^*$ | 1 | Maximum E2F activation rate |
| $k_R^*$ | 0.5 | Maximum REPAIR activation rate |
| $k_D^*$ | 1 | Maximum DSB generation rate |
| $k_B^*$ | 1 | Maximum 53BP1 activation rate |
| $d_E^*$ | 1 | E2F degradation rate constant |
| $d_R^*$ | 0.7 | REPAIR degradation rate constant |
| $d_B^*$ | 0.7 | 53BP1 degradation rate constant |
| $K_{GF,E}^*$ | 1 | Half-maximal constant for E2F activation by GF |
| $K_{E,E}^*$ | 0.09 | Half-maximal constant for E2F activation by E2F |
| $K_{GF,R}^*$ | 1 | Half-maximal constant for repair activation by GF |
| $K_{E,D}^*$ | 1 | Half-maximal constant for DSB generation by E2F |
| $K_{D,B}^*$ | 0.7 | Half-maximal constant for 53BP1 activation by DSB |
| $K_{GF,B}^*$ | 1 | Half-maximal constant for 53BP1 activation by GF |
| $K_{rep}^*$ | 1 | Half-maximal constant for DSB repair |
| $K_{inh}$ | 0.5 | Half-maximal constant for E2F inhibition by DSB |
| $k_{rep}$ | 2 | Maximum DSB repair rate |
| GF | 1, 0 | 1: before starvation or after restimulation, 0: during starvation |
| n | 6 | Hill coefficient for E2F and 53BP1 activation |
| $\tau_{GF \rightarrow E}$ | 2 | Time delay from GF to E2F |
| $\tau_{E \rightarrow E}$ | 0.5 | Time delay from E2F to E2F |
| $\tau_{D \rightarrow E}$ | 0.5 | Time delay from DSB to E2F |
| $\tau_{E \rightarrow D}$ | 0.5 | Time delay from E2F to DSB |

Initial condition: $[E2F] = [REPAIR] = [53BP1] = 0$ a.u.; $[DSB] = 4$ a.u.
Note 1: All parameters are defined in arbitrary units (a.u.), as the model is not explicitly parameterized to experimental/physical scales.
\* These parameters were doubled or halved for parameter sensitivity assay shown in Fig. S5.

